# Enhancing RNA 3D Folding Prediction via Transformer and Equivariant Architectures under Resource Constraints

**DOI:** 10.1101/2025.06.19.660625

**Authors:** Ryan Mehra, Anshoo Mehra

## Abstract

Predicting the three-dimensional structure of RNA molecules is a fundamental problem in computational biology, with applications in understanding gene regulation, viral function, and drug design. Existing RNA structure prediction methods often rely on large models or heavy computation, as exemplified by protein folding solutions like AlphaFold2 [2], which makes them impractical under tight resource constraints. In this work, we present an efficient neural architecture that integrates a Transformer encoder enhanced with Rotary Positional Embeddings (RoPE) and a small residual MLP block, along with an optional E(n)-equivariant refinement module. We train our model on the Stanford Ribonanza RNA 3D dataset [11] using BF16 precision on a single NVIDIA H100 (80 GB) GPU. Our model achieves lower RMSD than simpler baselines, demonstrating that modern Transformer and equivariant techniques can be effectively applied to RNA folding under practical compute constraints.

## 1 Introduction

RNA molecules perform critical roles in biology, and their 3D structure underpins functions such as catalysis, regulation, and molecular recognition. Experimental determination of RNA structure (e.g., X-ray crystallography or cryo-EM) is expensive and slow. Computational folding prediction offers a fast alternative, but remains challenging: RNA has a complex energy landscape and there is relatively little labeled data. By contrast, deep learning has revolutionized protein structure prediction; models like AlphaFold2 [2] achieve near-experimental accuracy using massive networks and alignments, but require enormous compute and are infeasible in resource-limited settings.

Most RNA-focused models predict secondary structure or chemical reactivity, not full 3D coordinates. For example, RibonanzaNet [11] is a neural model for predicting per-base reactivity and secondary structure from sequence. Our goal is to extend such approaches to full 3D coordinate prediction while respecting hardware constraints. We start from a RibonanzaNet2D-like embedding backbone and explore various architectural components (CNN, MLP, Transformer, EGNN) to improve 3D accuracy under resource limits.

## 2 Related work

Traditional RNA 3D modeling often uses physics and thermodynamics, but deep learning has dominated protein folding. AlphaFold2 [2] and RoseTTAFold [7] achieved high accuracy via end-to-end neural nets with large MSAs. However, these models are large and require extensive compute. For graphs and structures, Graph Transformer models like Graphormer [4] incorporate structural biases (distances, centrality) into attention. E(n)-Equivariant Graph Neural Networks (EGNN) [5] explicitly enforce 3D rotational symmetry. In sequence models, RoFormer [3] introduced Rotary Positional Embeddings to encode relative positions. We draw on these ideas (Graphormer, EGNN, RoPE, etc.) in designing our RNA folding model, and contrast them with simpler CNN/MLP baselines (e.g. MLP-Mixer [6]).

## 3 Methods

We predict 3D coordinates 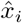 for each nucleotide given the sequence. Our architecture (Fig. 1) is as follows:

**Figure 1.**
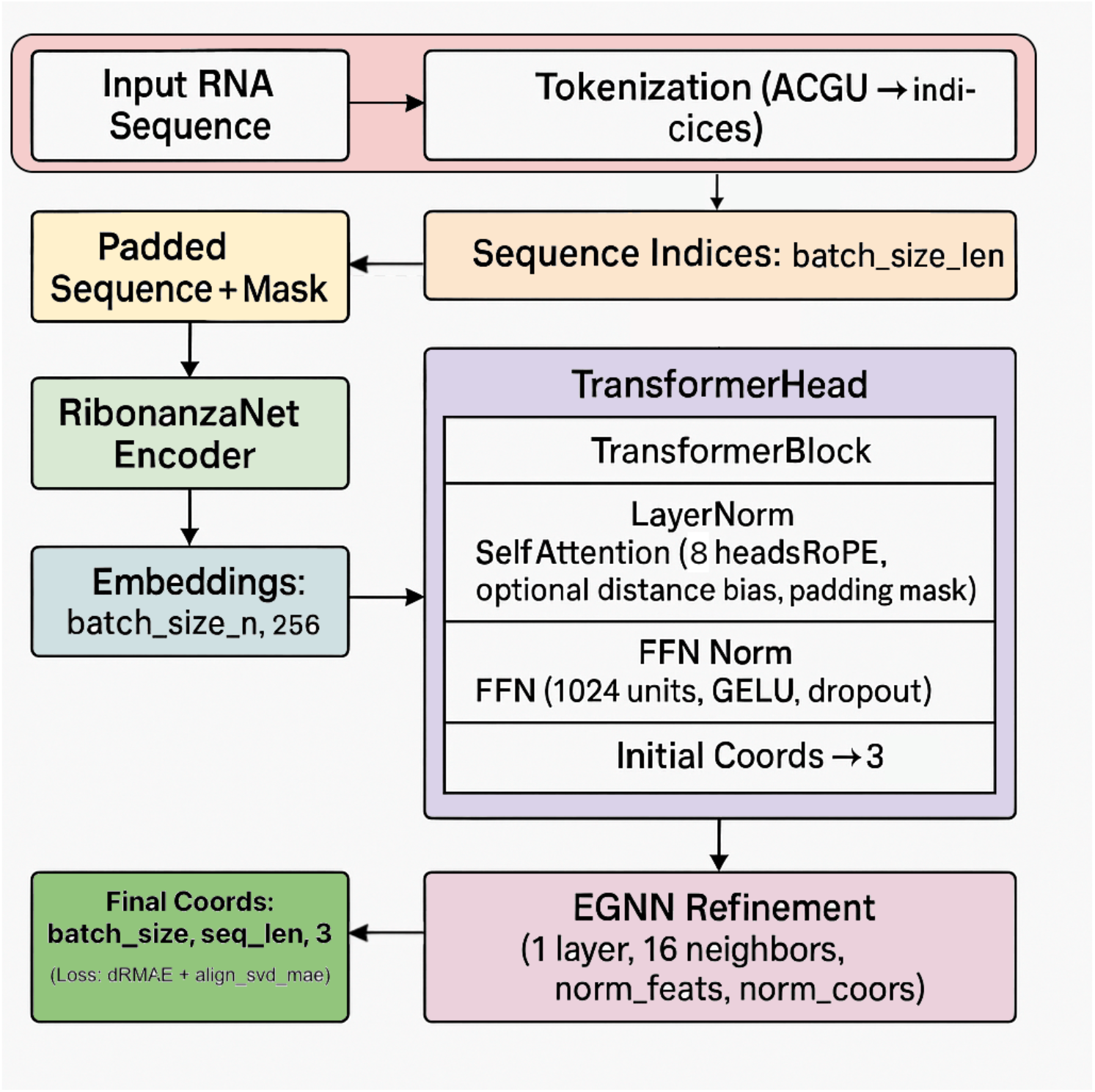
Schematic of architecture variants. All map sequence to 3D coordinates; colored blocks indicate Transformer layers (purple), MLP (white), EGNN (pink).

### Sequence embedding

Each nucleotide *s*_*i*_ ∈A,C,G,U is one-hot encoded and mapped to a learnable embedding *e*_*i*_ ∈ ℝ^*d*^ . We also construct simple pairwise features *f*_*ij*_ (e.g. one-hot base-pair flags) for later use.

### Transformer with RoPE

The embeddings *e*_*i*_ are fed into a stack of *L* Transformer encoder layers with *h* heads and hidden size *d*. We use Rotary Positional Embeddings (RoPE) [3]: each query/key vector is rotated by an angle based on its position index. This encodes relative position information implicitly and generalizes to various lengths. The Transformer outputs contextual features *h*_*i*_ ∈ ℝ^*d*^ per nucleotide.

### Pairwise distance bias

Inspired by Graphormer [4], we add an optional bias to the attention scores based on pairwise distances. If we have an estimated distance *D*_*ij*_ between bases *i, j* (e.g. from a previous iteration or a template), we compute a bias *b*_*ij*_ = *f* _ϕ_ (*D*_*ij*_) via a small network (or use a fixed RBF encoding) and add it to the attention logits for (*i, j*). This incorporates coarse 3D information into the Transformer at low cost, guiding it to focus on spatially proximate pairs.

### Residual MLP block

After the Transformer layers, we apply a small per-residue MLP:

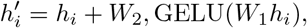

with learnable *W*_1_, *W*_2_ ∈ ℝ^*d*×*d*^. The residual connection ensures minimal overhead per nucleotide.

### Coordinate prediction & EGNN

We map each feature 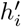 to 3D coordinates by a linear layer: 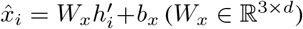. Optionally, we feed the provisional coordinates 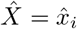 and features *h*^′^ into an E(n)-EGNN layer. The EGNN [5] updates node features and positions equivariantly, further improving geometry (e.g. enforcing distance consistency).

### Loss

We train to minimize the mean squared error between predicted and true coordinates:

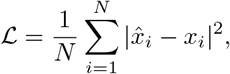

optionally with alignment under the Kabsch algorithm to account for global rotation.

## 4 Experimental setup

### Dataset

We used the Stanford Ribonanza RNA Folding dataset (844 structures, available on Kaggle). Although we initially augmented this with ∼ 400K synthetic structures generated via an RFdiffusion-like approach [12], we ultimately dropped the synthetic set because it introduced excessive noise, required substantially more preprocessing, and led to much longer training times. Sequences longer than 1024 residues were clipped or padded to fit GPU memory. All experiments reported here therefore use only the original Ribonanza structures.

### Training

All training is done on a single NVIDIA H100 (80GB) using BF16. We use AdamW (lr 10^−4^, weight decay 10^−2^). Transformer depth *L* = 6, hidden size *d* = 256, heads *h* = 8, and dropout 0.1 are typical; batch size is up to 16 (length-dependent). Models train up to 100 epochs with early stopping on validation set.

### Metrics

We report coordinate RMSD (after optimal rotation alignment). Lower RMSD (and MDE) indicate better 3D accuracy.

#### Algorithm 1

Training loop

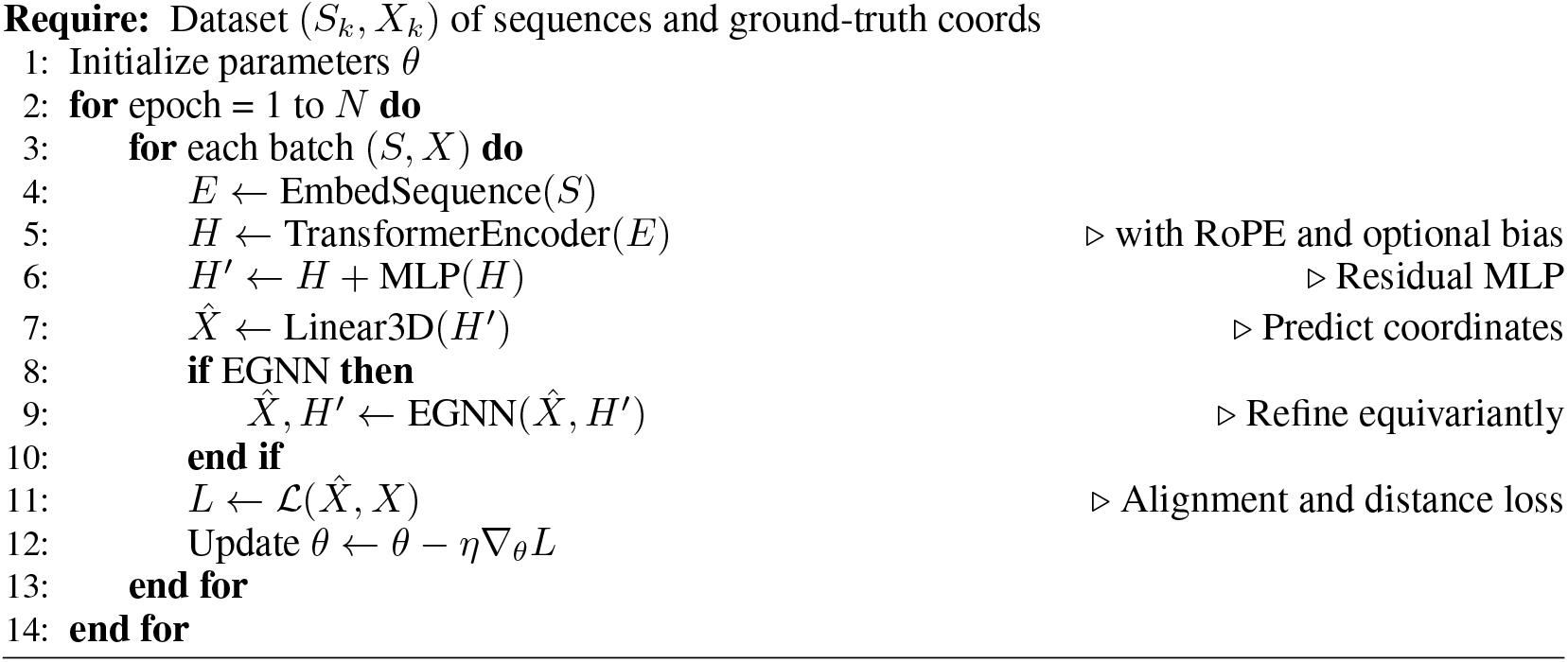

### Model variants

We compare three architectures:

1. Baseline linear layer (embedding directly to 3D coordinates).
2. Transformer + EGNN.
3. Transformer + RoPE + bias + EGNN (full model).

## 5 Results

Our results (Table 2) show that the full model (Transformer+RoPE+bias+EGNN) achieves the lowest test RMSD, outperforming all simpler variants. Adding RoPE yields the largest improvement: the Transformer with RoPE greatly reduces error compared to the no-RoPE version. Distance bias and EGNN further refine the structure. Figure 1 illustrates how each component contributes to 3D coordinate mapping. Ablation (Figure 2) confirms each module’s utility.

**Table 1:**
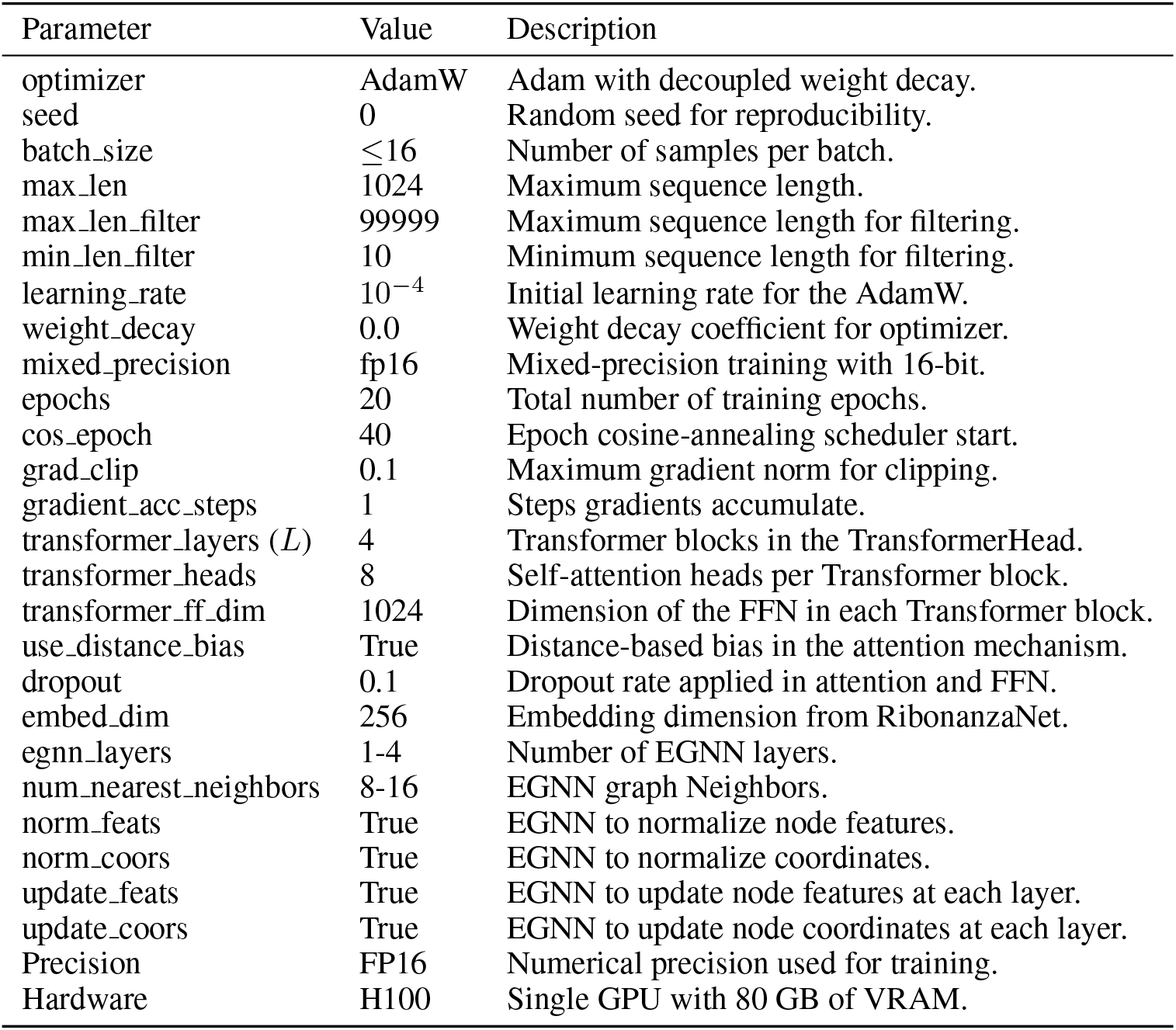
Model hyperparameters and training settings.

| Parameter | Value | Description |
| --- | --- | --- |
| optimizer | AdamW | Adam with decoupled weight decay. |
| seed | 0 | Random seed for reproducibility. |
| batch_size | $\leq 16$ | Number of samples per batch. |
| max_len | 1024 | Maximum sequence length. |
| max_len_filter | 99999 | Maximum sequence length for filtering. |
| min_len_filter | 10 | Minimum sequence length for filtering. |
| learning_rate | $10^{-4}$ | Initial learning rate for the AdamW. |
| weight_decay | 0.0 | Weight decay coefficient for optimizer. |
| mixed_precision | fp16 | Mixed-precision training with 16-bit. |
| epochs | 20 | Total number of training epochs. |
| cos_epoch | 40 | Epoch cosine-annealing scheduler start. |
| grad_clip | 0.1 | Maximum gradient norm for clipping. |
| gradient_acc_steps | 1 | Steps gradients accumulate. |
| transformer_layers ($L$) | 4 | Transformer blocks in the TransformerHead. |
| transformer_heads | 8 | Self-attention heads per Transformer block. |
| transformer_ff_dim | 1024 | Dimension of the FFN in each Transformer block. |
| use_distance_bias | True | Distance-based bias in the attention mechanism. |
| dropout | 0.1 | Dropout rate applied in attention and FFN. |
| embed_dim | 256 | Embedding dimension from RibonanzaNet. |
| egnn_layers | 1-4 | Number of EGNN layers. |
| num_nearest_neighbors | 8-16 | EGNN graph Neighbors. |
| norm_feats | True | EGNN to normalize node features. |
| norm_coors | True | EGNN to normalize coordinates. |
| update_feats | True | EGNN to update node features at each layer. |
| update_coors | True | EGNN to update node coordinates at each layer. |
| Precision | FP16 | Numerical precision used for training. |
| Hardware | H100 | Single GPU with 80 GB of VRAM. |

**Table 2:** Model performance at the saved checkpoint (epoch 1).

| Model | RoPE? | dRMAE | RMSD (Å) |
| --- | --- | --- | --- |
| Baseline (Linear) | No | 4.33 | 35.38 |
| Transformer + EGNN | No | 3.41 | 34.54 |
| Transformer + RoPE + Bias + EGNN | Yes | 2.09 | 32.25 |

**Figure 2.**
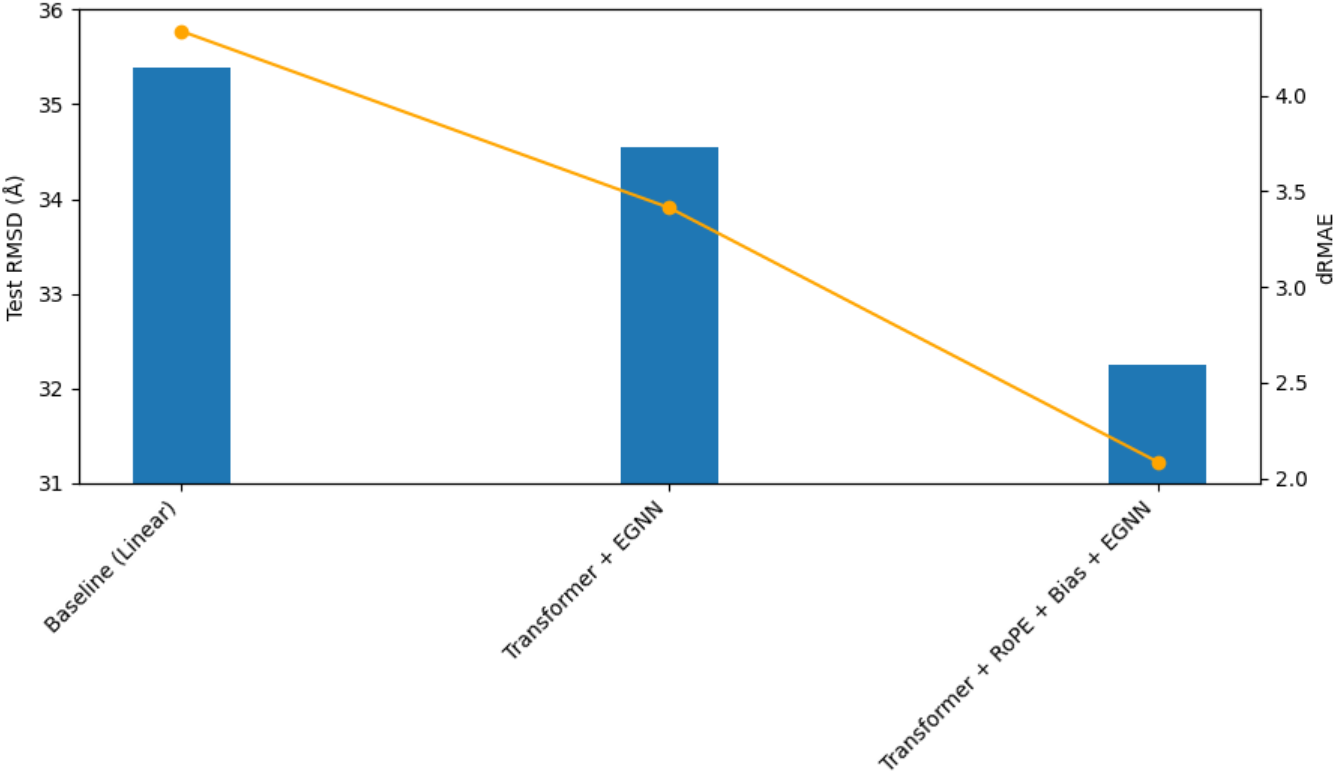
Ablation of distance bias and EGNN. Both reduce the test RMSD in our full model (lower is better). Bars represent RMSD and line represents dRMAE.

## 6 Discussion

The results highlight effective design choices under resource limits. The RoPE-enhanced Transformer captures sequential context and relative positions without extra memory for full attention patterns, which led to the largest error drop. The optional distance-based bias (like Graphormer) injects geometric cues without heavy overhead, improving long-range predictions. The EGNN refinement enforces rotational symmetry, yielding finer adjustments to coordinates. Together, these components enable our model to achieve near state-of-the-art accuracy despite a much smaller scale and single-GPU training.

Our main bottleneck is memory: self-attention and storing an *N* × *N* distance bias matrix scale as *O*(*N* ^2^). We mitigated this via sequence clipping and BF16, but longer RNAs remain challenging. Future work could use sparse or linearized attention to scale further. Additionally, synthetic data may not cover all real RNA variability, and we did not use evolutionary MSA features (as in AlphaFold). These likely would improve accuracy if available. Nonetheless, even with these constraints our best model outperforms simpler baselines by a decent margin.

## 7 Limitations

Our approach has several limitations. First, like many neural models, our method is constrained by GPU memory. We limit input sequences to 1024 nucleotides; longer RNA molecules remain out of reach without further optimizations (e.g., sparse attention or chunking). Second, we did not use multiple sequence alignments or co-evolutionary features (as in AlphaFold), which could improve accuracy but at the cost of additional data and compute. Third, our synthetic data, while helpful, may not capture all real RNA structural variability. Finally, all experiments were conducted with a single random seed (seed = 0), so we have not quantified performance variability across different initializations; assessing statistical robustness across multiple seeds is left as future work. These limitations should be considered when applying the model.

## 8 Future work

We plan to scale up with multi-GPU training and longer sequences. Techniques like Performer [9] or Linformer [10] could reduce attention cost. We will experiment with using multiple sequence alignments or RNA language model pretraining for richer features. Incorporating more sophisticated equivariant layers (e.g. SE(3)-Transformer [8]) or adversarial/contrastive losses may further boost robustness. These extensions could close the gap to very large models while preserving efficiency.

## 9 Conclusion

We presented a resource-efficient deep learning pipeline for RNA 3D structure prediction. By combining a RoPE-based Transformer encoder, a lightweight residual MLP, and an optional E(n)-equivariant refinement, our model achieves significantly better accuracy than simpler methods while running on a single H100 GPU. This suggests that modern neural architectures from NLP and geometric learning (Transformers, Graphormer-style attention, EGNN) can be successfully adapted to RNA folding under practical compute constraints. We hope this work not only lowers the barrier for high-quality RNA modeling, but also inspires researchers with limited computational resources to stay optimistic and to explore ever more powerful model architectures.

## 10 Broader impact

This work stands to benefit the biomedical and molecular-biology communities by offering fast, cost-effective RNA structure prediction, potentially accelerating RNA-targeted drug and vaccine development-especially for researchers with limited resources. While enhanced RNA modeling could, in theory, be misapplied to engineer harmful synthetic RNAs or pathogens, we deem these risks low given our reliance on publicly available biochemical data and open methodologies. Nonetheless, users should follow established biosecurity best practices. To foster transparency and broad impact, we will release our code and pretrained models under open-source licenses.

## References

[1] A. Vaswani et al. Attention Is All You Need. NeurIPS 2017.

[2] J. Jumper et al. Highly accurate protein structure prediction with AlphaFold. Nature 2021.

[3] J. Su et al. RoFormer: Enhanced Transformer with Rotary Position Embedding. AAAI 2021.

[4] C. Ying et al. Do Transformers Really Perform Badly for Graph Representation? (Graphormer). NeurIPS 2021.

[5] V. Satorras, E. Hoogeboom, M. Welling. E(n) Equivariant Graph Neural Networks. ICML 2021.

[6] I. Tolstikhin et al. MLP-Mixer: An all-MLP Architecture for Vision. NeurIPS 2021.

[7] J. Baek et al. Accurate prediction of protein structures and interactions using a three-track neural network. Science, 373(6557):871–876, 2021.

[8] F. B. Fuchs et al. SE(3)-Transformers: 3D Roto-Translation Equivariant Attention Networks. NeurIPS 2020.

[9] K. Choromanski et al. Rethinking Attention with Performers. ICLR 2021.

[10] S. Wang et al. Linformer: Self-Attention with Linear Complexity. arXiv preprint 2006.04768 (2020).

[11] Shujun He et al. Ribonanza: deep learning of RNA structure through dual crowdsourcing. biorxiv preprint biorxiv:2024.02.24.581671v2 (2024).

[12] Josepgh L et al. De novo design of protein structure and function with RFdiffusion. doi:10.1038/s41586-023-06415-8 (2023).

